# PLI Analyzer for Data-driven Validation of AI Predicted Biomolecular Interfaces

**DOI:** 10.1101/2025.09.29.678845

**Authors:** Frances Liang, Suhas Srinivasan, Howard Y. Chang

## Abstract

PLI-Analyzer is a computational tool for detailed analysis of protein, RNA, and DNA interaction interfaces, combining atomic-level contact detection with domain annotations from UniProt and InterPro. It offers customizable interaction thresholds, residue-level output, and compatibility with AI-generated structures, enabling precise validation and interpretation of contact dynamics. We applied PLI-Analyzer to evaluate high-quality protein-protein complexes predicted by AlphaFold-3 and Boltz-2, revealing notable differences in predicted interfaces and confidence scores. These findings highlight current limitations in generative structure models specifically for macromolecular complexes and underscore the need for robust, domain-informed evaluation frameworks in structural bioinformatics.

## Introduction

Recent advancements in structural prediction tools such as AlphaFold are becoming an essential tool in the discovery of the biological mechanisms and functions of protein, RNA, and DNA complexes [1] [2]. Proteins do not function in isolation, as their biological roles are realized through intricate interactions with other proteins, RNA, and DNA, forming dynamic molecular complexes that drive cellular processes. This understanding is central to how we investigate molecular structures, as the ability to predict and model these interactions is critical to accurately capturing the true biological function of these molecules [3]. These prediction tools are now transforming how we investigate molecular structures, offering valuable mechanistic insights that were previously accessible only through expensive and time-consuming experimental approaches [4]. Consequently, structure prediction has become a key tool in molecular biology, drug discovery, and synthetic biology [2].

These methodological advancements in protein structure prediction have enabled the creation of large-scale structural resources like the AlphaFold Protein Structure Database, which contains over 200 million protein structures of almost all proteins known to science [5]. These models can generate multiple structures from a single sequence, each with subtle variations that may reflect uncertainties in specific regions or potential alternate conformational states. Additionally, the accuracy of current models generally diminishes when transitioning from monomers to multimers [1], due in part to the combinatorial complexity of multimeric interfaces and the relative scarcity of high-resolution, experimentally determined multimeric structures for training. For these reasons, reliable means of evaluation and validation of these predictions are essential. Specifically, robust evaluation based on established structural features and known protein annotations is therefore critical for selecting the most biologically relevant models [6]. Without robust methods to evaluate predicted structures, researchers risk basing downstream analyses, mutagenesis experiments or therapeutic designs on incorrect structural predictions.

To address this need, we developed the tool PLI-Analyzer (Protein ligand interaction-Analyzer, PLIA), which evaluates and scores multimer predictions—from any method—based on the degree to which predicted structures overlap with known interacting sites along the protein sequence. PLI-Analyzer is a computational tool designed to analyze interaction interfaces in protein, RNA, and DNA complexes (https://github.com/SuhasSrinivasan/plia). It first identifies contact regions through atomic interactions based on Voronoi diagrams [7] and then compares these interaction sites to known interacting motifs from UniProt and InterPro’s domain annotations. The tool supports customization of interaction thresholds and residue padding and offers intermediate output generation at the residue level to enable detailed, stepwise analysis of interacting contacts and regions. This feature allows for deeper inspection of interaction patterns, making it easier to trace, interpret, and validate contact dynamics throughout the analysis pipeline. Users can input structures generated from AI methods, and the tool provides the detected interacting regions, optionally aligned with UniProt’s known interacting regions. This tool is particularly useful for validating predicted interfaces and characterizing novel interaction sites in biomolecular complexes.

We used the tool to evaluate the accuracy of two state-of-the-art structure prediction methods, namely AlphaFold-3 (AF3) and the recent Boltz-2 (BZ2) [8] method for 30 randomly sampled known protein-protein interactors. First, known protein-protein interaction domains were mined from UniProt and InterPro to create a high-quality dataset of human proteins with interaction sites. This curated data was used to randomly sample binary interactions from the human HINT database (High-quality INTeractomes) [9]. The sequences for these binary interactors were submitted to the AlphaFold Server, and Boltz-2 for obtaining the 3D predicted structures of the binary PPI complexes.

Each protein complex was analyzed through PLI-Analyzer to compare predicted interaction sites with known interacting domains annotated in UniProt and InterPro. Predicted sites were filtered to include only motifs composed of at least three consecutive residues. Each filtered site was then compared to known domains using sequence identity. For identity scoring, the highest score obtained against any known domain was recorded as the representative match.

## Results

Our analysis of the 30 predicted PPI complexes included multiple metrics and revealed that AF3 and BZ2 models had average sequence identities of 61.2% and 56.2%, respectively, when compared to known interacting regions (Fig. 1B). Average sequence identity in the case of AF3 ranged from a minimum of 26% to a maximum of 95% with a standard deviation of 19% (Supp. Fig. 8A). Average sequence identity for BZ2 ranged between 21% and 97% with a standard deviation of 19% (Supp. Fig. 8B). Additionally, for both models, most identity scores for each interactor were in the 20–60% or 80–100% range, with very few regions falling in the 0–20% or 60–80% range (Supp. Fig. 7). Specifically, the lack of predictions in the 0–20% range suggests that both AF3 and BZ2 can correctly identify at least a portion of the interacting regions.

**Figure 1:**
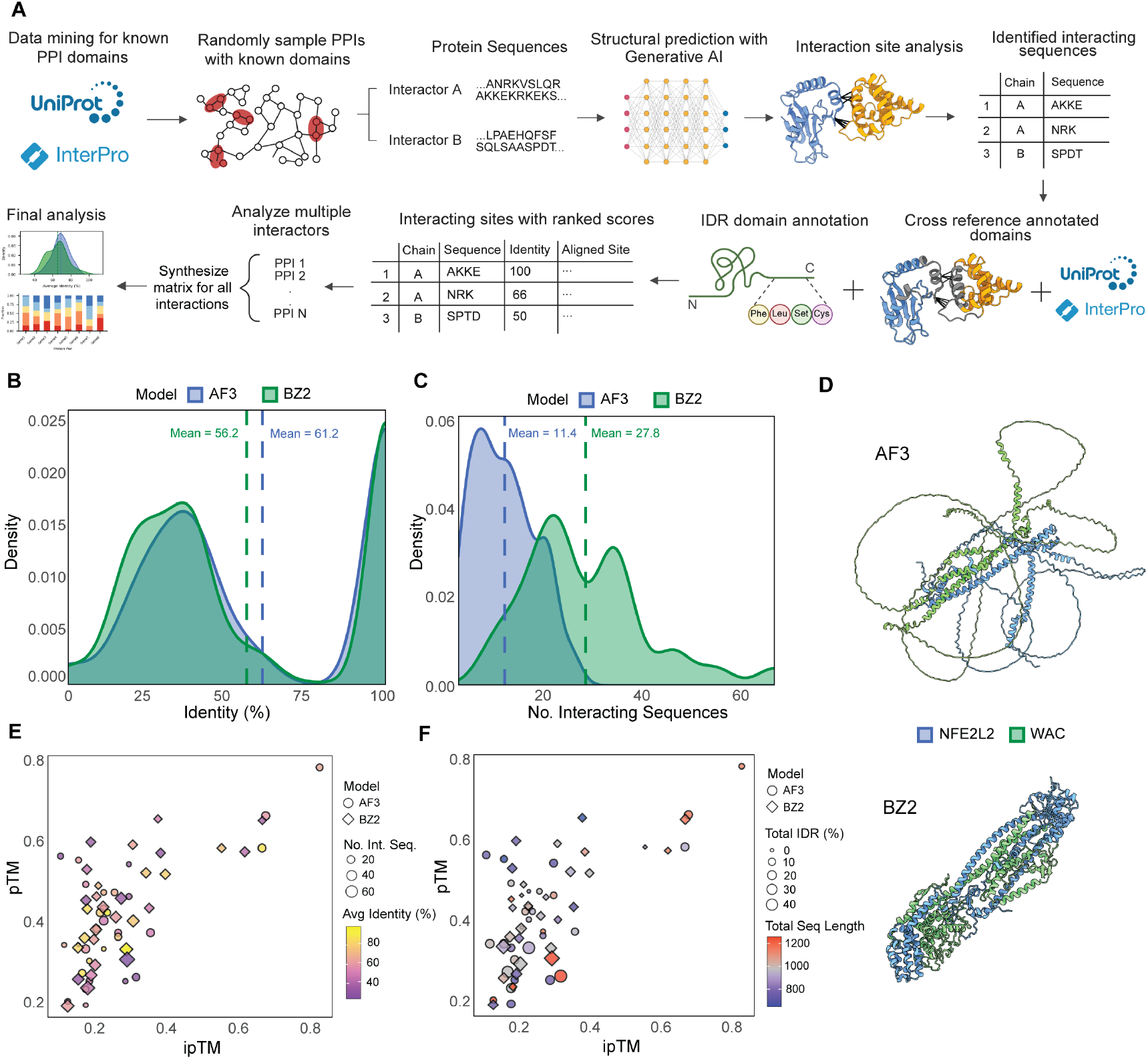
Analytical workflow for PLI-Analyzer and evaluation of state-of-the-art AI structure prediction methods. **A**, Data curation and analysis workflow of PLI-Analyzer for 30 randomly sampled high-quality human binary interactions. **B**, Distribution of sequence identity between predicted and known interaction sites for AF3 and BZ2. **C**, Distribution of the number of predicted interacting sites across AF3 and BZ2. **D**, AF3 and BZ2 predicted protein structures for protein interactor pair NFE2L2 (blue) and WAC (green). **E**, Scatterplot of, pTM vs ipTM, number of interacting sequences (size), avg. identity (color), for AF3 (circle) and BZ2 (diamond). **F**, Scatterplot of pTM vs ipTM, total IDR percentage (size), total sequence length (color), for AF3 (circle) and BZ2 (diamond).

One of the most significant differences between AF3 and BZ2 is in the average number of predicted interacting sequences per complex (Fig. 1C). While AF3 predicted 11.4 interacting sequences on average, BZ2 predicted 27.8, almost three times as many. An example of this difference can be seen in the interactor pair NFE2L2 and WAC (Fig. 1D). AlphaFold generated 16 interacting sites for this complex, whereas Boltz-2 predicted 66. Upon further analysis, Boltz-2 predicted a highly packed and organized structure, while AlphaFold generated multiple long stretches of intrinsically disordered regions (IDRs). Cross-referencing with UniProt regional annotations confirmed that WAC has 444 residues defined as IDRs, which accounted for approximately 68% of the total protein length. Similarly, ∼30% of NFE2L2 is also annotated as disordered. Thus, the BZ2 prediction strongly misrepresents the NFE2L2-WAC complex as it predicts little to no disordered regions. While AF3’s prediction characterized significantly more of the structure as disordered, over 25% of these predicted disordered regions were not annotated as IDRs. Together, this was evidence of large disparities between previously defined structural properties and predicted structures, particularly within the context of IDRs, with BZ2’s predicted number of interacting sequences having greater correlation with TotaI IDR of the protein complex and specifically IDR of Protein B (Supp. Fig. 3B and Supp. Fig. 6B).

pTM (predicted TM-score) and ipTM (interface predicted TM-score) are two of the main confidence metrics for structural prediction algorithms like AF3 and BZ2. pTM estimates the global structural accuracy of a predicted protein complex, with scores above 0.5 suggesting a possibly correct overall fold. ipTM assesses the confidence in the relative positioning of subunits within a complex, where values above 0.8 indicate high-quality interface predictions, and values below 0.6 suggest likely failure. The relatively low average pTM and ipTM scores for both AF3 (pTM: 0.41, ipTM: 0.269) and BZ2 (pTM: 0.443, ipTM: 0.283) suggested that neither model consistently produced highly confident structural predictions across the evaluated protein complexes (Supp. Fig. 4). Boltz-2’s slightly higher pTM score indicates better internal consistency in overall structure, but its lower ipTM score indicates lower confidence in the predicted interaction sites themselves, which was reflected in slightly decreased interface prediction identity (Supp. Fig. 5).

We also see a strong correlation between ipTM and pTM scores in both AF3 and BZ2 (Supp. Fig. 6). However, BZ2 also demonstrated a strong negative correlation between pTM and the number of predicted interacting sequences, implying that as the number of predicted interacting sites for a pair of proteins increased, the model’s confidence in the accuracy of individual structures tended to decrease (Supp. Fig. 6B).

For AF3, ipTM scores displayed a moderately positive correlation with the number of interacting sequences, while BZ2 demonstrated the opposite with a moderately negative correlation. In contrast, BZ2’s ipTM were relatively stable regardless of protein size or identity. Notably, pTM scores for both models were largely consistent across the full range of protein sizes (Fig. 1F).

## Discussion

As structural prediction tools like AlphaFold-3 and Boltz-2 become increasingly integral to molecular biology and structural biology, evaluating the biological relevance of their predictions for multimeric complexes is more critical than ever. With the explosion of predicted structural data, tools such as PLI-Analyzer create a much-needed framework to assess whether predicted interaction sites align with established functional annotations, helping researchers avoid drawing conclusions that contradict known characteristics. Our evaluation across 30 high-quality binary protein complexes revealed that while both AlphaFold-3 and Boltz-2 could identify interaction interfaces with moderate accuracy, their performance varies significantly depending on the context, particularly in the presence of IDRs. For instance, the over structuring of binary interactor NFE2L2 and WAC artificially inflates the number of predicted interacting sequences by misclassifying disordered regions as potential interaction sites, leading to predictions that would fail to represent the complex’s functional reality.

While recent work, such as Predictomes, has made valuable contributions to understanding PPIs, their approach filters for high-confidence (pLDDT ≥ 50) structures, and relies solely on resolved structures from the PDB for training data, thereby restricting the types of interactions that can be analyzed [10], and our approach compliments this technique by bringing in annotation information of protein domains and regions and can handle low-complexity regions (IDRs).

Ultimately, our work further shines a light on current limitations of generative structures of proteins and biomolecular complexes from the latest AI methods and therefore emphasizes the need for further investment by the structural bioinformatics community to develop comprehensive and robust evaluation techniques to critically assess generated structures using multi-modal domain knowledge and functional data. We hope that tools like PLI-Analyzer might be one of the tools to help bridge the gap between raw structural predictions and biologically meaningful insights.

## Notes

https://github.com/SuhasSrinivasan/plia

### Competing Interest Statement

The authors have declared no competing interest.

