## Supplement for "PLI Analyzer for Data-driven Validation of AI Predicted Biomolecular Interfaces"

### *Supplementary Analysis, Figures and Methods*

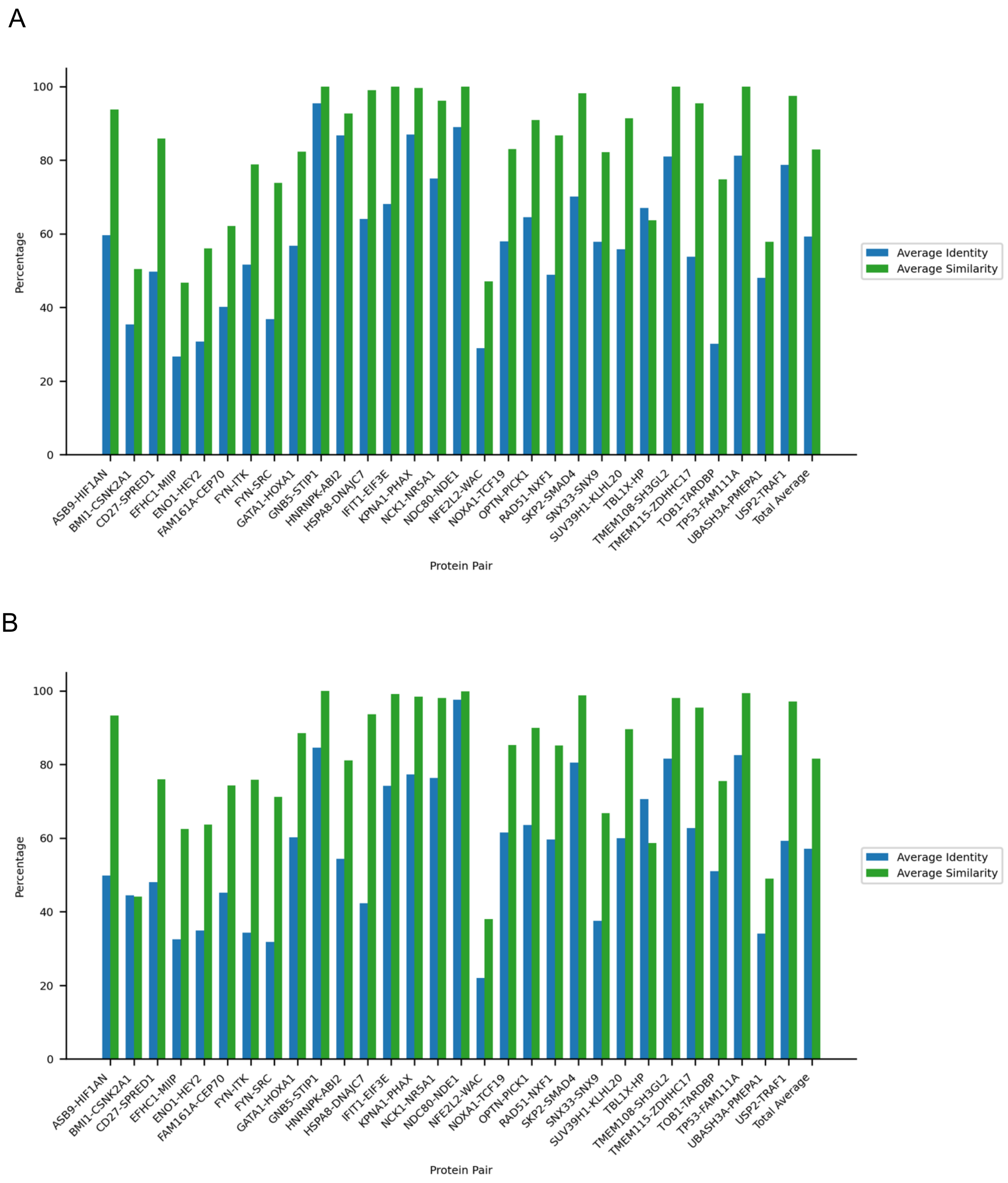


**Supplementary Figure 1**: Bar graphs of identity and similarity scores for predicted binary interactors of AF3 (A) and BZ2 (B).

A

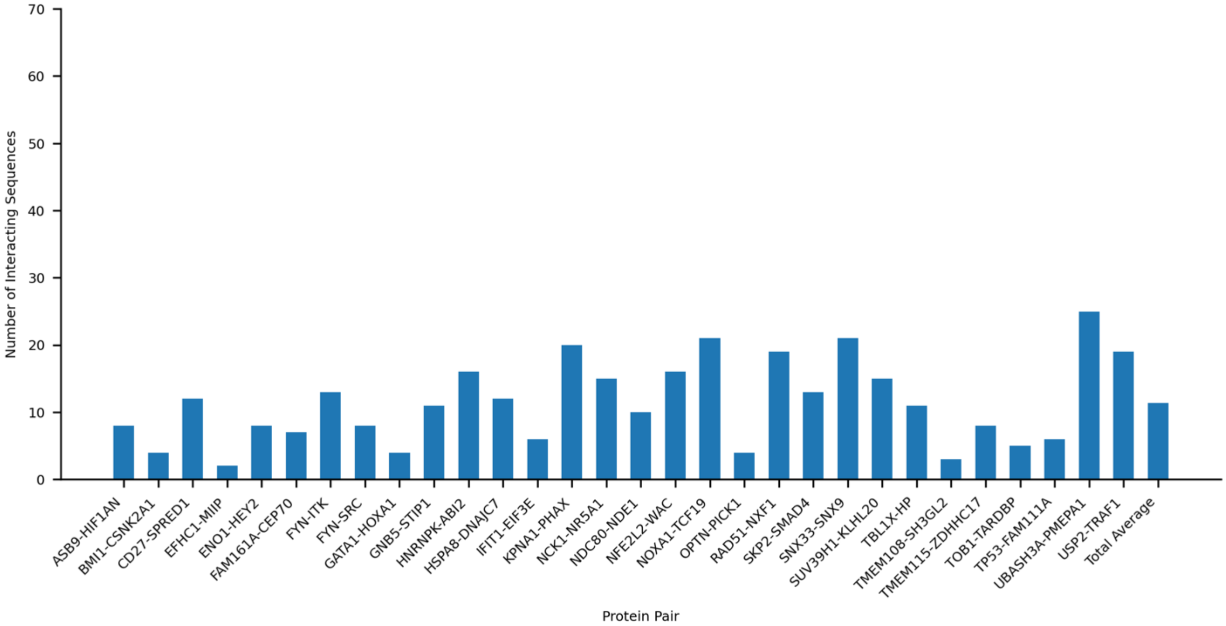


B

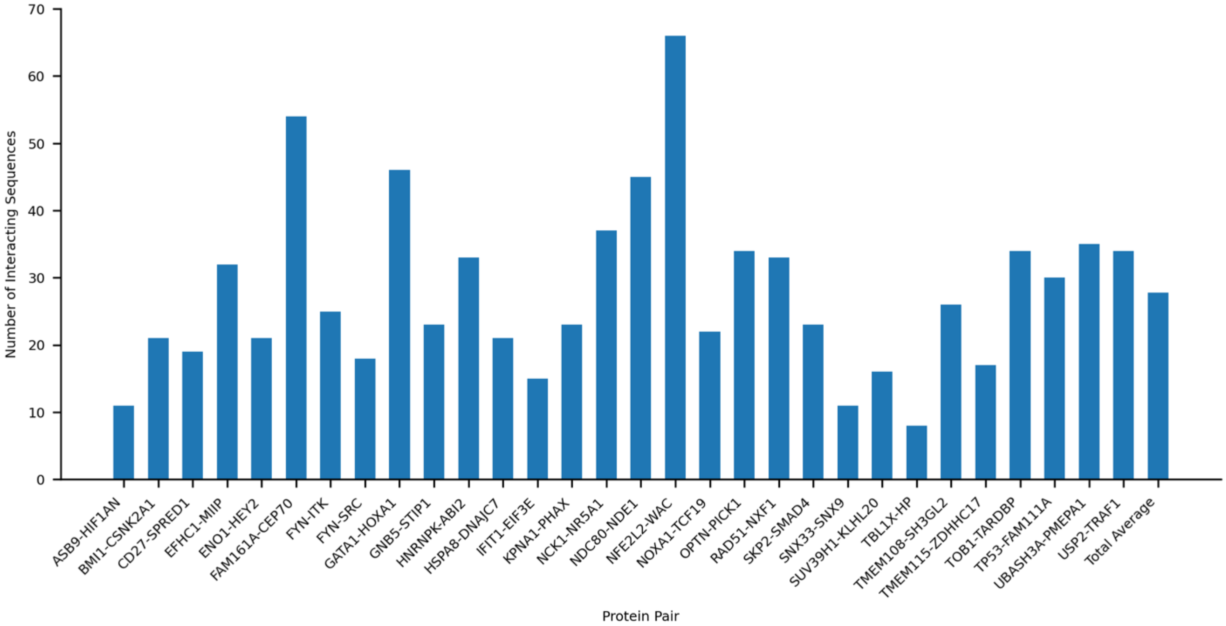


**Supplementary Figure 2**: Bar graphs of the number of predicted interaction sites across AF3 (A) and BZ2 (B).

A

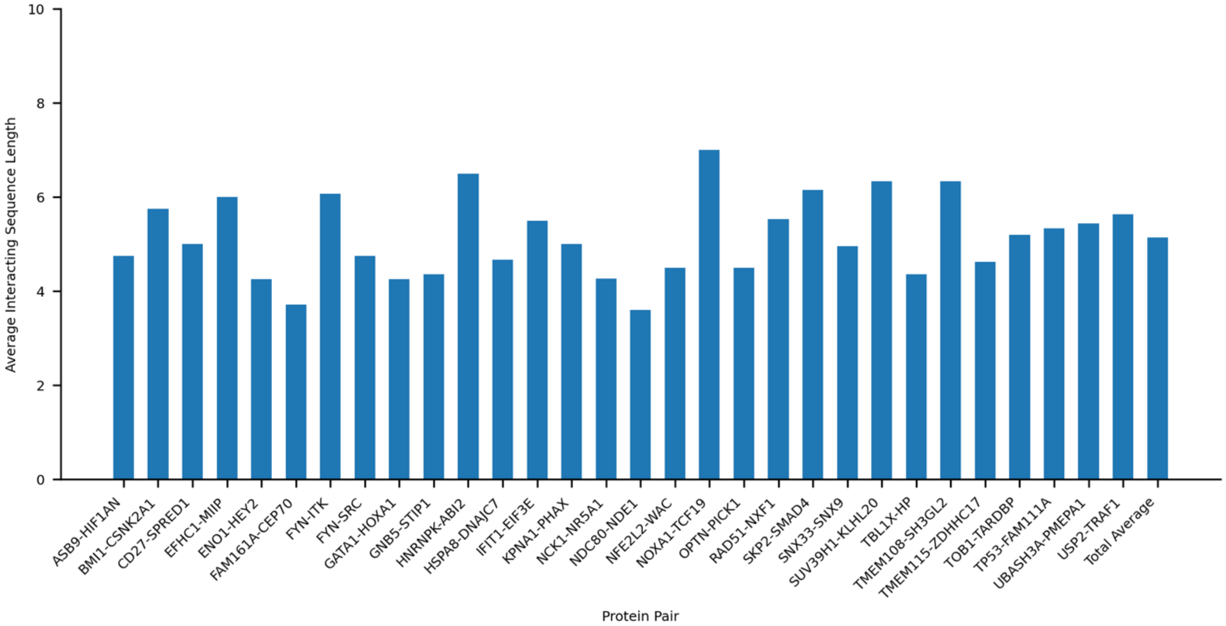


B

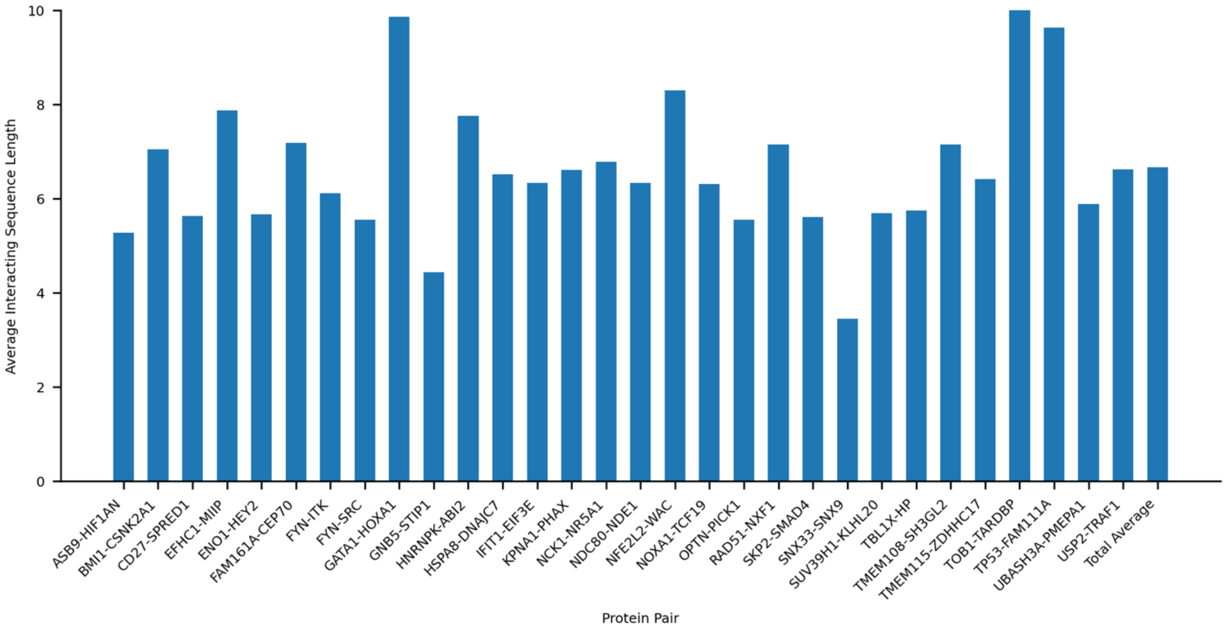


**Supplementary Figure 3**: Bar graphs of the lengths (number of residues) of predicted interaction sites in AF3 (A) and BZ2 (B).

A

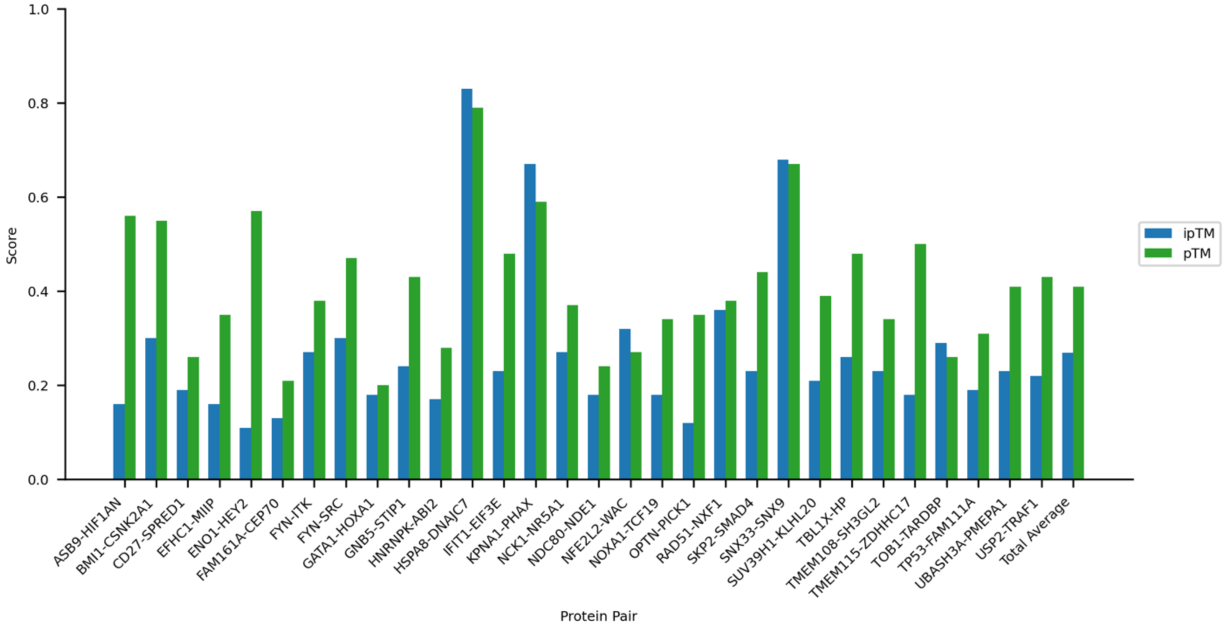


B

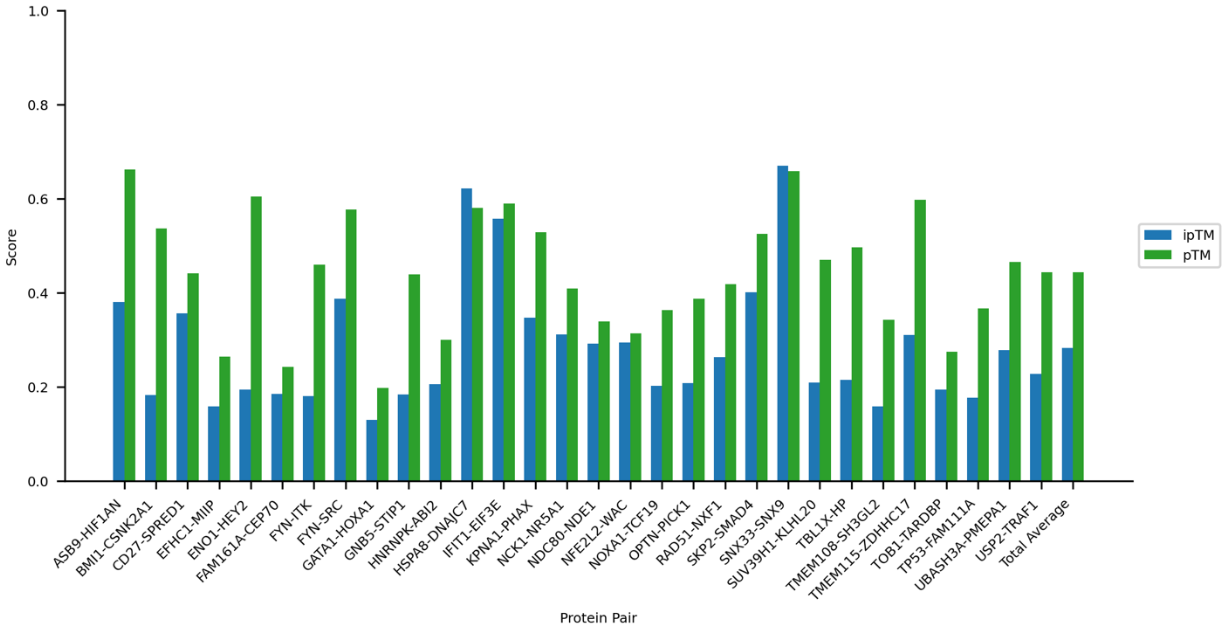


**Supplementary Figure 4**: ipTM and pTM structural confidence scores for predicted protein interactors of AF3 (A) and BZ2 (B).

A

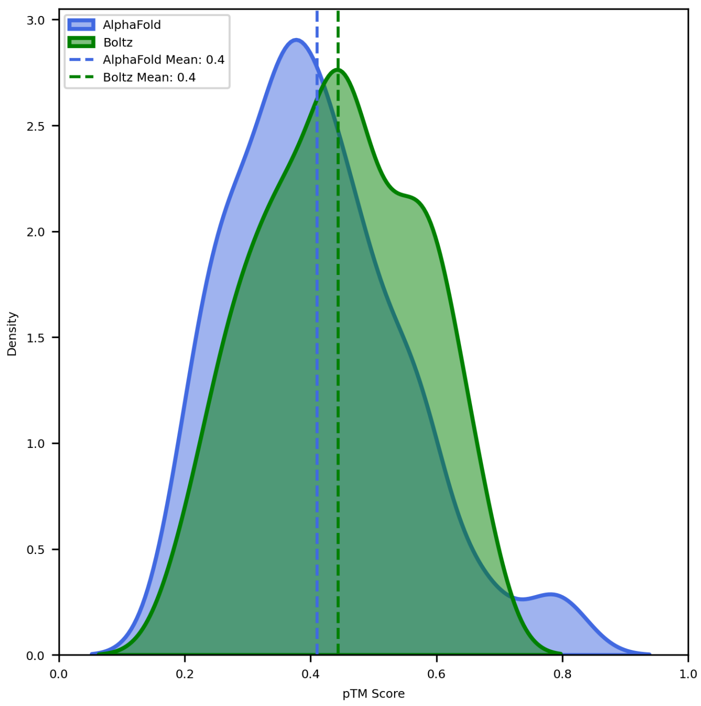


B

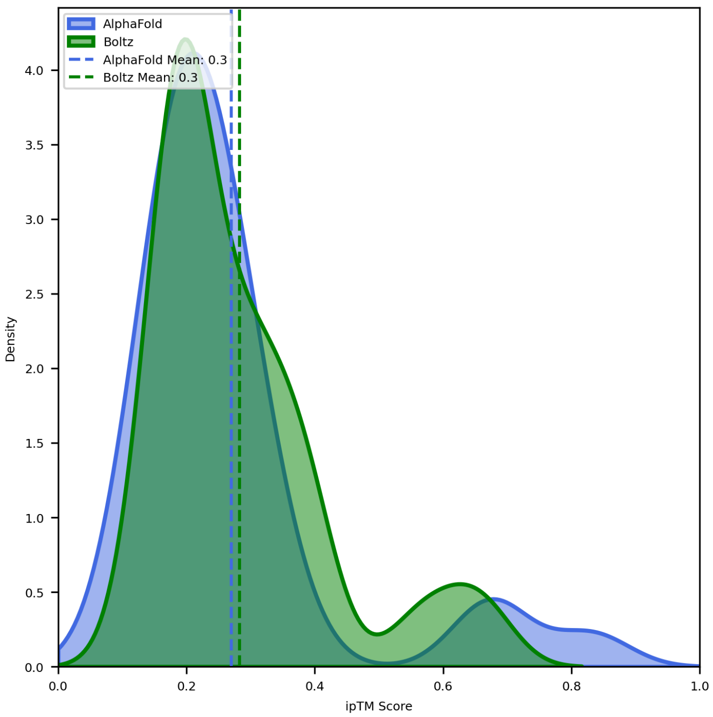


**Supplementary Figure 5**: Density estimates of pTM (A) and ipTM (B) scores for AF3 and BZ2 predictions.

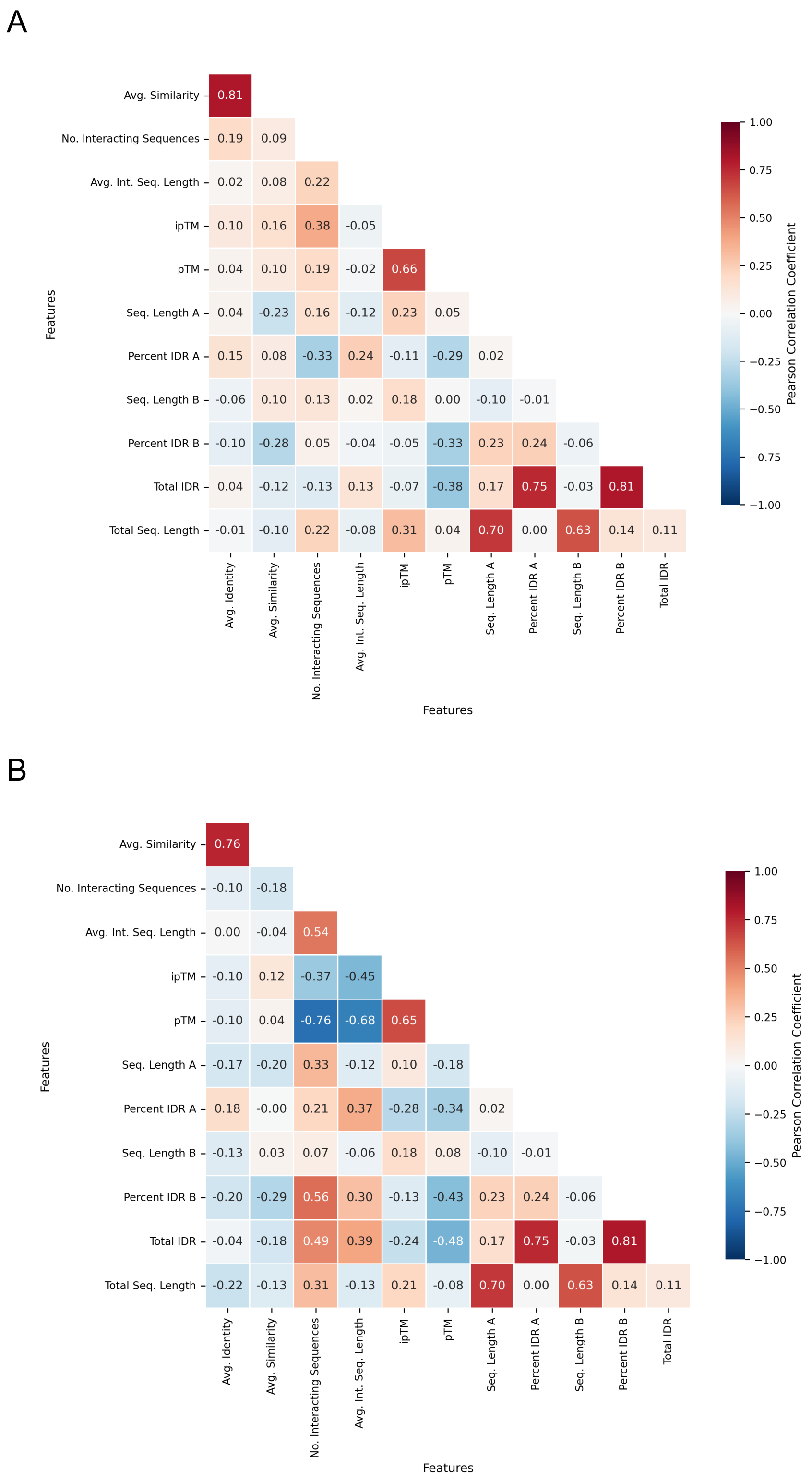


**Supplementary Figure 6**: Pearson correlation heatmaps of interaction confidence scores and interaction associated attributes for AF3 (A) and BZ2 (B).

A

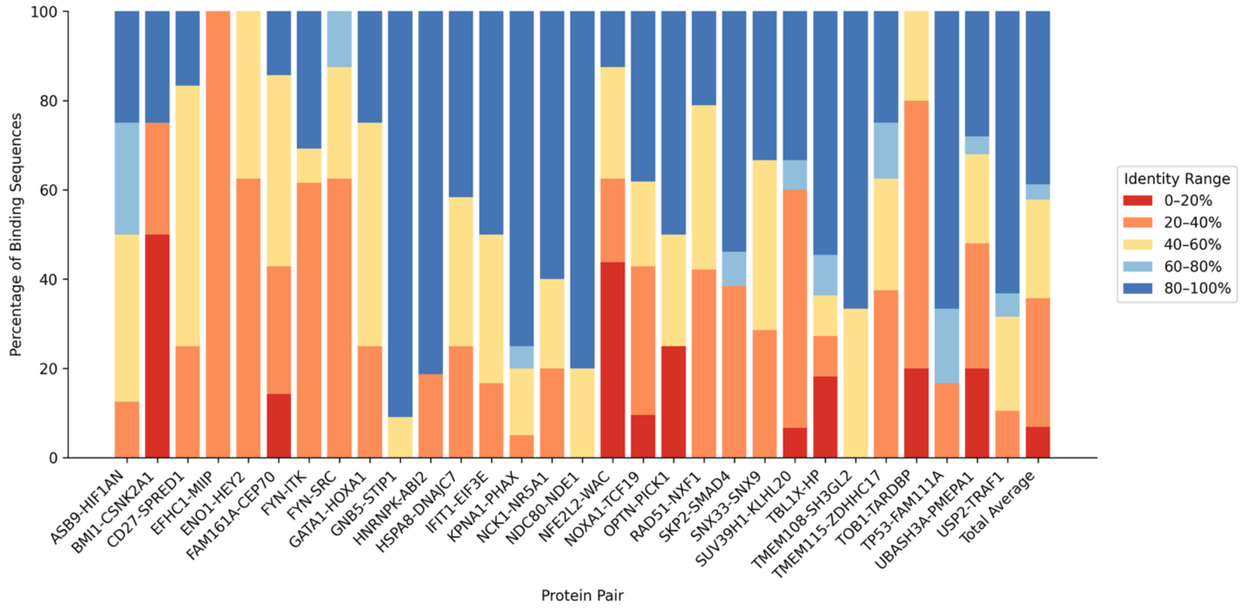


B

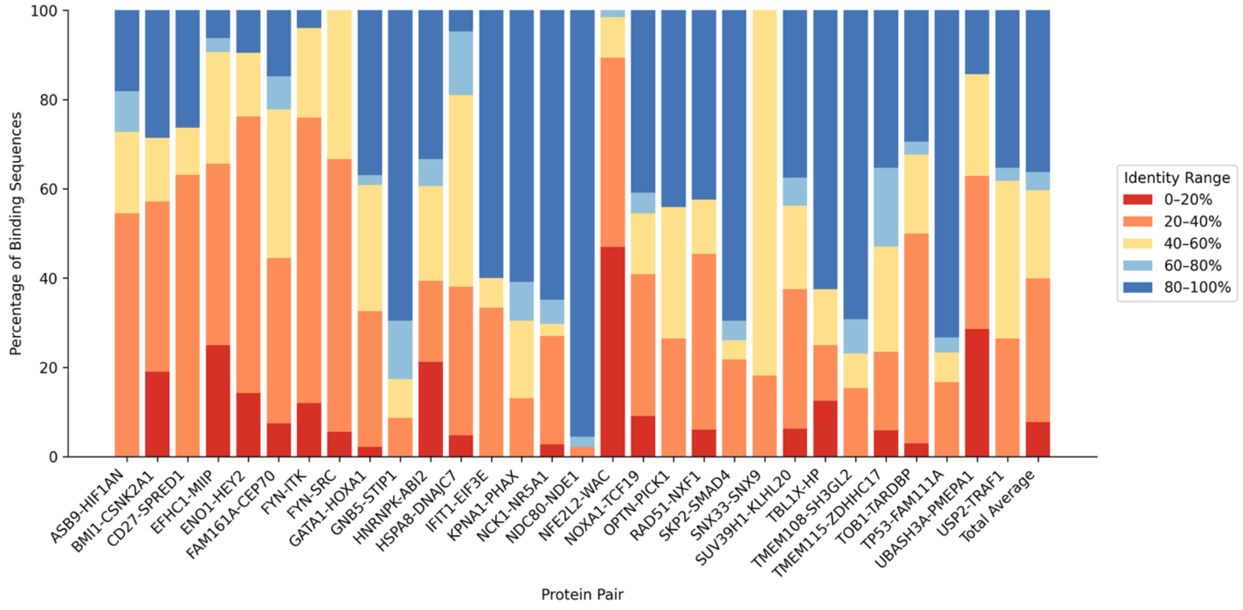


**Supplementary Figure 7**: Stacked bar plots displaying distribution of sequence identity ranges across AF3 (A) and BZ2 (B).

A

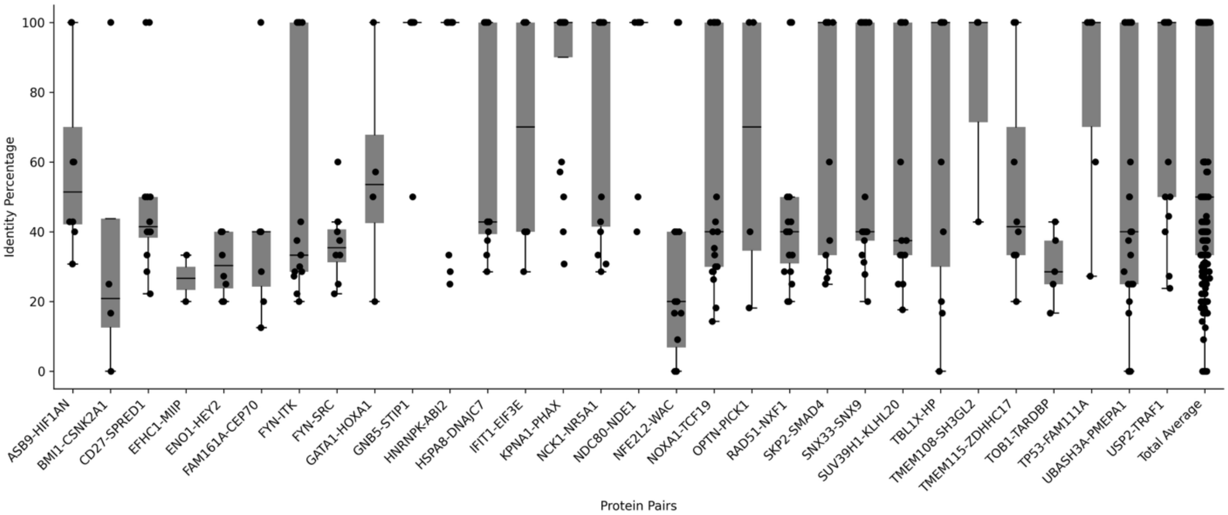


B

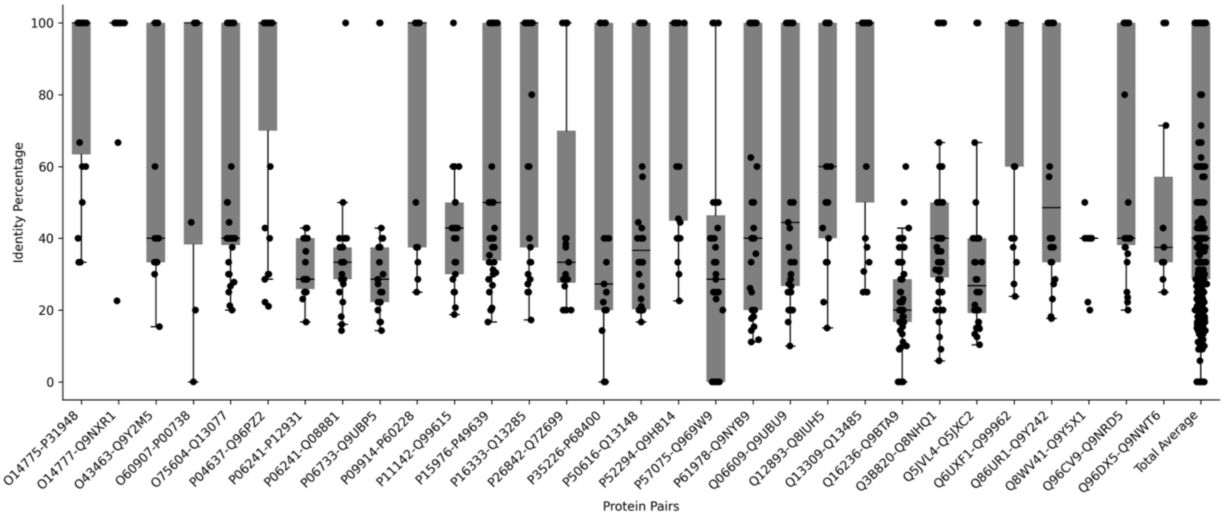


**Supplementary Figure 8**: Boxplot representing sequence identities across the binary interactors for AF3 (A) and BZ2 (B).

### Methods

The Protein–Ligand Interaction Analyzer (PLIA) is a modular structural bioinformatics workflow for identifying, extracting, and evaluating macromolecular interaction interfaces—primarily protein-protein interactions—at residue resolution. PLIA was designed for high-throughput validation and characterization of interfaces in experimentally determined or AI-predicted assemblies from methods such as AlphaFold 3, Boltz 2 and RosettaFold All-Atom [1-3]. Typical applications include: (i) verifying that predicted complex interfaces recover known binding motifs or domains with high identity or similarity, (ii) prioritizing predicted complexes whose interfaces show strong concordance to curated references, and (iii) and surveying large structural datasets for trends in interface motif recovery (e.g., identity/similarity distributions, segment lengths) that may guide experimental follow-up.

The workflow emphasizes (i) robust geometric detection of inter-chain contacts in protein complexes, (ii) principled assembly of contiguous, sequence-level interface segments with configurable padding, and (iii) quantitative comparison of extracted interface segments to curated reference annotations of binding motifs and interaction domains derived from UniProt and InterPro. PLIA operates on PDB and mmCIF structural inputs, integrates Voronoi-based contact geometry via Voronoi diagrams [4], and uses Biopython [5]. When available, per-structure confidence metadata from AlphaFold is summarized alongside the interface assessments to contextualize results.

#### *Input data model and preprocessing*

PLIA accepts experimentally determined assemblies or predicted models in PDB (or mmCIF through conversion noted below) format [6]. Structures are parsed to extract the first model and enumerate unique inter-chain pairs within that model. Water and other solvent entities are excluded from contact analysis. Non-polypeptide residues (e.g., nucleic acids, ligands) are detected to avoid conflating their atoms with protein contacts during protein–protein interface extraction; interface scoring, as described below, is currently defined for polypeptide chains.

To analyze each input complex, PLIA requires a per-interactor reference sequence provided by the user either through new experiments such as cross-linking mass spectrometry [7] or published literature [8] or queried from large-scale community resources, such as UniProt [9] and InterPro [10], containing residue spans corresponding to known interaction motifs or domains. We provide tools to obtain UniProt annotation when working with structures derived from standard UniProt sequences. These spans serve as sequence-level targets for evaluating extracted interface segments. By decoupling structural parsing from reference knowledge, PLIA remains agnostic to the origin of structures while leveraging widely curated biological annotations.

For maximal compatibility we also provide a tool to convert mmCIF inputs files to PDB format while preserving chain identifiers and residue numbering [6]. Only standard amino acids are considered when deriving final segment sequences; non-standard residues are ignored for the purposes of sequence comparison.

#### *Geometric contact detection*

PLIA's interface identification uses Voronoi tessellation [4] of atomic balls as implemented in Voronota [11]. In this formalism, each atom is represented as a sphere with an associated radius; Voronoi decomposition assigns spatial regions to atoms, and inter-atomic contacts correspond to shared Voronoi faces. Compared with fixed distance cutoffs, Voronoi-face contacts better account for atomic radii and the geometry of local packing, improving precision at complex interfaces [].

The contact detection stage proceeds as follows:

1. Atomic-ball generation and inter-atomic contact computation are performed for the full complex. Solvent contacts and intra-chain contacts are excluded, ensuring that only inter-chain contacts contribute to interface identification.
2. A minimum shared Voronoi-face area threshold is applied to retain physically meaningful contacts while suppressing incidental near approaches [12].
3. Optionally, a distance-based pruning can be applied downstream to remove residue pairs whose minimal inter-atomic distance exceeds a user-specified cutoff (Å). This dual-criterion approach (area-based primary detection, optional distance pruning) balances sensitivity with specificity across diverse structural contexts.
4. Contacts are deduplicated to unique residue–residue pairs, and only contacts involving standard amino acids are retained for sequence assembly.

#### *Residue-to-segment assembly with contextual padding*

The residue-level contact list is reduced to minimal contiguous sequence segments that define interface footprint(s) on each chain:

1. For each chain in a contacting pair, PLIA maps atom-level contacts to residue indices and merges consecutive residue indices into contiguous runs (e.g., 37–42, 265–275).
2. Since functionally relevant interaction motifs frequently extend beyond residues with direct atomic contacts, PLIA expands each contiguous run by a configurable padding (default: ±1 residue) on both termini, clipping to valid chain bounds. Padding helps stabilize motif-level comparisons by including immediate sequence context [13].
3. Standard amino-acid residues are converted to one-letter codes for downstream scoring. Very short segments are removed using a minimal length threshold (default: 3 residues) to reduce noise from sparse or marginal contacts.

#### *Quantitative sequence evaluation against reference motifs*

Extracted interface segments are quantitatively compared to the per-interacting chain reference annotations. The aim is to capture both exact local matches (identity) and conserved, gap-permitting similarities (similarity) at the motif scale, rather than enforcing whole-domain global alignments [14, 15].

Sequence identity (ungapped) is computed as the fraction of exact amino-acid matches when the shorter segment is slid across the longer reference span without gaps, reporting the best local frame, formally defined as below:

$$Sequence\ Identity = \frac{Number\ of\ identical\ positions}{Total\ number\ of\ aligned\ positions} \times 100$$

This gap-free, local framing emphasizes strict residue-level concordance within the motif window and is robust to segment length differences.

Sequence similarity is computed from a local, gap-permitting pairwise alignment optimized for maximal match count under a simple scoring scheme and normalized by the query length. This measure tolerates small sequence variations (insertions/deletions) that frequently occur at interface loops and motif boundaries, complementing the stricter ungapped identity. Reporting both identity and similarity separates strict positional conservation from conservative, indel-tolerant similarity.

For each interface segment, PLIA evaluates all reference spans and reports the best-matching reference along with its identity and similarity scores. This procedure yields, per chain, a set of scored segment–reference relationships that are ranked and summarized.
